# Metabolic signatures of cardiorenal dysfunction in plasma from sickle cell patients, as a function of therapeutic transfusion and hydroxyurea treatment

**DOI:** 10.1101/2023.04.05.535693

**Authors:** Angelo D’Alessandro, S. Mehdi Nouraie, Yingze Zhang, Francesca Cendali, Fabia Gamboni, Julie A. Reisz, Xu Zhang, Kyle W. Bartsch, Matthew D. Galbraith, Joaquin M. Espinosa, Victor R. Gordeuk, Mark T Gladwin

## Abstract

Metabolomics studies in sickle cell disease (SCD) have been so far limited to tens of samples, owing to technical and experimental limitations. To overcome these limitations, we performed plasma metabolomics analyses on 596 samples from patients with sickle cell sickle cell disease (SCD) enrolled in the WALK-PHaSST study. Clinical covariates informed the biological interpretation of metabolomics data, including genotypes (hemoglobin SS, hemoglobin SC), history of recent transfusion (HbA%), response to hydroxyurea treatment (HbF%). We investigated metabolic correlates to the degree of hemolysis, cardiorenal function, as determined by tricuspid regurgitation velocity (TRV), estimated glomerular filtration rate (eGFR), and overall hazard ratio (unadjusted or adjusted by age). Recent transfusion events or hydroxyurea treatment were associated with elevation in plasma free fatty acids and decreases in acyl-carnitines, urate, kynurenine, indoles, carboxylic acids, and glycine- or taurine-conjugated bile acids. High levels of these metabolites, along with low levels of plasma S1P and L-arginine were identified as top markers of hemolysis, cardiorenal function (TRV, eGFR), and overall hazard ratio. We thus uploaded all omics and clinical data on a novel online portal that we used to identify a potential mechanism of dysregulated red cell S1P synthesis and export as a contributor to the more severe clinical manifestations in patients with the SS genotype compared to SC. In conclusion, plasma metabolic signatures – including low S1P, arginine and elevated kynurenine, acyl-carnitines and bile acids - are associated with clinical manifestation and therapeutic efficacy in SCD patients, suggesting new avenues for metabolic interventions in this patient population.

## INTRODUCTION

Sickle cell disease (SCD) encompasses a group of inherited hemoglobinopathies caused by mutations in the gene coding for the beta globin chain (*HBB*) in the hemoglobin tetramer.^1,2^ These mutations result in the substitution of single amino acid residue, E6 to V in sickle Hb (HbS) allele β^S^, or E6 to K in HbC. Under deoxygenated conditions, sickling of red blood cells (RBCs) occurs following crystallization of mutant globins into macrofibers,^3-5^ which in turn disrupts erythrocyte morphology, rheology^6^ and metabolism.^7^ Clinically, these molecular events manifest themselves in a complex disease characterized by hemolytic anemia, inflammation and recurrent vaso-occlusive episodes.^8^ While such manifestations tend to be more severe in patients with the SS genotype, vaso-occlusive complications are still present in patients with the SC genotype.^2,9^

Lifelong transfusion represent a mainstay in the management of SCD in children and adults.^8^ Increases in circulating levels of heme and iron secondary to the elevated hemolytic propensity of sickle red blood cells (RBCs) triggers recurrent episodes of vaso-occlusion and inflammation, which progressively cause damage to most organs, including the kidneys, lungs, and the cardiovascular system, and overall reduced life expectancy.^8^ Interventions based on treatment with hydroxyurea (hydroxycarbamide) have shown a certain degree of efficacy in up-regulating the expression of fetal hemoglobin (HbF) and increasing mean corpuscular volume (MCV),^10^ while decreasing the frequency of pain episodes and other acute complications in adults and children with sickle cell anemia of SS or Sβºthal genotypes, reducing the incidence of life-threatening neurological events such as primary stroke by maintaining transcranial Doppler velocities.^11^

The circulating metabolome offers a window not just on chemical individuality under healthy conditions^12^, but also onto dysregulation of systems homeostasis under pathological conditions, such as in the case of SCD.^7^ Studies on the circulating metabolome in SCD have identified potential new avenues for metabolic interventions. For example, arginine supplementation^13^ has been proposed as a strategy to boost nitric oxide synthesis and NO-dependent vasodilation, thus counteracting hemoglobin-dependent NO scavenging secondary to intravascular hemolysis, and release of red cell arginase which has been shown to catabolize L-arginine.^13,14^ Similarly, hemolysis-derived free hemoglobin, heme and iron have been associated with increased renal, cardiopulmonary and infectious complications in this and other patient populations.^15,16^ An alternative metabolic intervention. glutamine supplementation has been proposed as a strategy to boost glutathione synthesis, thus counteracting oxidant stress in the sickle RBC.^17,18^ More recently, metabolomics interventions have been designed to boost the rate of late, payoff steps enzymes of glycolysis such as pyruvate kinase.

This strategy promotes the synthesis of adenosine triphosphate and energy metabolism in the sickle cell, while depleting the reservoirs of allosteric modulators like 2,3-diphosphoglycerate that promote sickling by stabilizing the tense deoxygenated state of hemoglobin.^19^ Other metabolomics studies have identified dysregulated adenosine signaling^20,21^ and sphingolipid metabolism^22^ as potential targets for therapeutic interventions, though studies have not progressed beyond promising pre-clinical animal models. One major limitations of almost all metabolomics studies of SCD plasma to date is that limited sample size and lack of extensively curated clinical data hampered the extrapolation of the translational relevance of the reported findings. To bridge this gap, here we performed a metabolomics study of plasma from 596 SCD patients, for which extensive information on patients’ demographics (sex, age, BMI), genotypes, hematological and clinical characteristics, as well as therapeutic interventions (transfusion events, as inferred from HbA %; hydroxyurea treatment) were available. Results from analyses in plasma were compared against the recently characterized RBC metabolome in this cohort^23^, identifying compartment-specific metabolic signatures associated with potential novel strategies for metabolic interventions in SCD.

## METHODS

An extended version of this section is provided in **Supplementary Methods Extended**, and our previous metabolomics report on RBC samples from the same cohort.^23^

### Ethical statement, blood collection, processing and storage

A total of 596 SCD patients with available plasma samples were recruited as part of the Walk-PHaSST clinical trial (NCT00492531 at ClinicalTrials.gov and HLB01371616a BioLINCC study) under protocols approved by the University of Pittsburgh^9,24-28^. Clinical covariates were collected prior to plasma metabolomics analysis, including hemolysis, tricuspid regurgitation velocity (TRV),^29^ parameters indicative of kidney function such as estimated glomerular filtration rate (eGFR), creatinine.).^9,24-28^ Standard HPLC methods were used to determine the percentage of hemoglobin S (sickle - HbS), C (HbC), F (fetal - HbF) and A (alpha, normal - HbA).^9,24-28^

### Ultra-High-Pressure Liquid Chromatography-Mass Spectrometry (MS) metabolomics

Metabolomics extractions,^30,31^ analyses^32,33^ and elaborations^34^ were performed as previously described.

### Statistical analyses

Multivariate and multivariable analyses, including principal component analyses (PCA), hierarchical clustering analyses, two-way ANOVAs, correlation analyses (Spearman) and calculation of receiver operating characteristic (ROC) curves were performed using MetaboAnalyst 5.0.^35^ For survival analysis, we used time to right censorship (including death or last follow-up) as time to event and vital status (death or alive) was the studied outcome. PCA was used to derive a hemolytic component from lactate dehydrogenase, aspartate aminotransferase, total bilirubin and reticulocyte percent.^27,36,37^ We applied Cox proportional hazard models to calculate the hazard ratios and P value for each metabolite. Cox analysis and adjusted regressions of metabolites to clinical outcomes were performed in R (R Core Team (2022), https://www.r-project.org/).

### Sickle Cell Disease ShinyApp portal

A portal for online sharing of all the data generated in this study, based on code from the COVID-Ome Explorer portal.^38^

### Data sharing statement

All results described in this study are provided in **Supplementary Table 1** and accessible at https://mirages.shinyapps.io/SCD2023/.

## RESULTS

### Metabolomics of the WALK-PHASST SCD cohort

Metabolomics analyses were performed on plasma samples from 596 sickle cell study participants in the screening phase of Walk-PHASST cohort (**Figure 1.A**). Results extensively reported in **Supplementary Table 1**, which also includes anonymized patients’ biological data, including genotype, sex, age, body mass index (BMI – **Figure 1.A**). An unsupervised analysis of the plasma metabolomics results is provided in **Supplementary Figure 1**. Principal component analyses (PCA) and hierarchical clustering analyses (HCA) of all the metabolomics data showed a significant separation of plasma samples from individuals with SS genotypes compared to the other groups (**Supplementary Figure 1.A-B**). As previously noted,^23^ despite double randomization of the samples and blinding of the analytical team, plasma creatinine levels measured via mass spectrometry significantly correlated with clinical measurement for this metabolite (**Supplementary Figure 1.C**).

**Figure 1.**
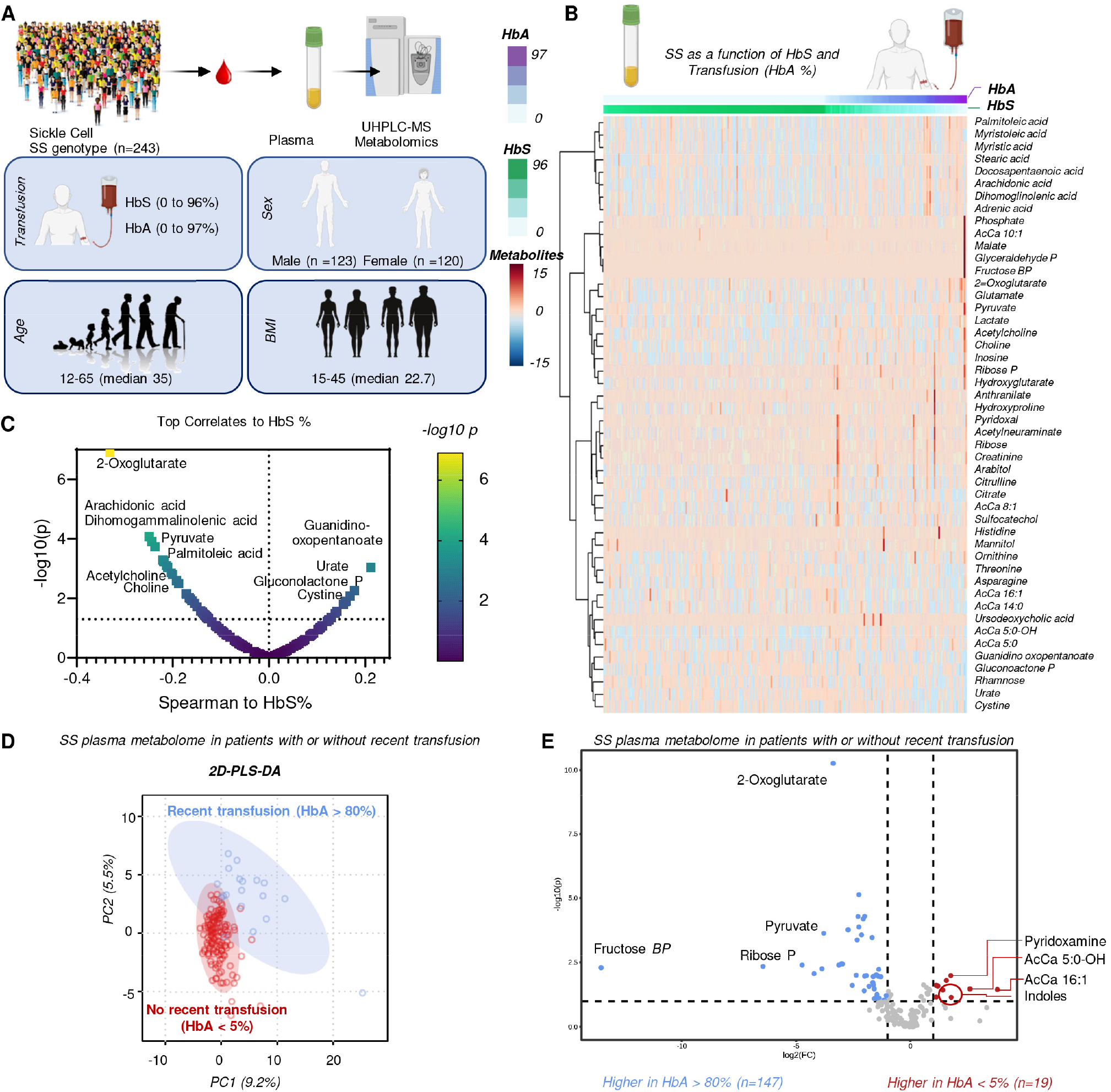
Plasma metabolomics of the WALK-PHASST SCD cohort. The plasma metabolome of patients with SS genotype – who did not receive other treatment (e.g., hydroxyurea) was evaluated as a function of recent transfusion events, as determined by HbA % (n=123 males and 120 females - **A**). In **B**, hierarchical clustering analysis of plasma metabolomics data as a function of HbS and HbA% (top 50 significant metabolites are shown). In **C**, smile plot of plasma metabolic correlates to HbS%. In **D**, partial least-square discriminant analysis of plasma metabolomics data in SS patients with recent (HbA>80%) or non-recent (HbA<5%) transfusion events. PC=Principal component. In **E**, volcano plot of plasma metabolites significantly higher in patients with no recent transfusion event (HbA <5%) or with recent exchange transfusion (HbA>80%). FC=fold-change.

### Plasma metabolic profiles in SS vs SC patients who have not been recently transfused

Individuals with SS genotypes represented ∼73% of the cohort in this study and showed distinct clinical^9,24-28^ and metabolic phenotypes (**Supplementary Figure 1**) compared to patients with other genotypes. As such, the first analysis we performed focused on the SS patient population. Clinically, many of these patients are routinely transfused, which could be herein assessed by screening of HbA percentages. We used this intervention to all for an in vivo assessment of the metabolome of plasma from patients with high levels of HbS compared with patients with SS genotype who have very high levels of transfused AA blood as a control. For this first analysis, other treatments (e.g., hydroxyurea – *see below*) were excluded. HCA of plasma metabolites significantly affected by HbA and HbS levels in SS patients identified a significant increase in the levels of free fatty acids (palmitoleic, myristoleic, myristic, stearic, docosapentaenoic, arachidonic, dihomogammalinolenic and adrenic acid) as a function of high levels of HbA, as opposed to declines in a series of acyl-carnitines (5:0, 5:0 OH, 14:0; 16:1 – **Figure 1.B**). Elevated HbS levels were correlated (Spearman; p<0.05) with increases in urate, gluconolactone phosphate (a metabolite of the oxidative phase of the pentose phosphate pathway – PPP) and cystine, with decreases in the levels of pyruvate, choline and acetylcholine, on top of the aforementioned free fatty acids (**Figure 1.C**). To further evaluate the impact of recent transfusion events on the plasma metabolome of SS patients, data were binned into extremes HbA phenotypes, with two groups identified based on “no recent transfusion events” (HbA<5%) or “recent transfusion events” (HbA>80%) that showed distinct metabolic phenotypes by partial least square-discriminant analysis (PLS-DA – **Figure 1.D**). Volcano plots of significant metabolites in these two groups further identified transfusion-associated elevation in pyruvate and other pentose or hexose phosphate compounds (fructose bisphosphate and ribose phosphate) and decreases in indoles, acyl-carnitines (especially those derived from branched chain amino acid catabolism – 5:0 and 5:0 OH) and pyridoxamine (**Figure 1.E**).

### Impact of hydroxyurea treatment on the plasma metabolome

Of the 433 patients with SS genotypes enrolled in this study, 306 had not been recently transfused (HbA <20%), of which 147 had received treatment with hydroxyurea (**Supplementary Figure 2.A**). Unsupervised analyses including PCA and HCA identified a modest impact of hydroxyurea treatment on the plasma metabolome, with the strongest impact on the levels of acyl-carnitines, urate, L-arginine and amino acids (lower in treated patients compared to untreated ones – **Supplementary Figure 2.B-C**). These effects were corroborated by correlation with HbF %, which increased after hydroxyurea treatment and was also positively associated with elevated tryptophan and indole metabolites (indole, indole acetate – **Supplementary Figure 2.D**). Linear models, unadjusted or adjusted by HbF% confirmed a strong impact of hydroxyurea on plasma L-arginine, acyl-carnitines and amino acids (**Supplementary Figure 2.E**). Acknowledging the limited signal emerged from this analysis, perhaps as a result of uncertain adherence to the hydroxyurea regimen, we performed an additional analysis by focusing on SS patients on hydroxyurea with mean corpuscular volumes (MCV) > 105 (**Supplementary Figure 2.F**), owing to the well-established effect on this hematological parameter of hydroxyurea treatments.^10^ This analysis unmasked a signal related with hydroxyurea-dependent decrease in circulating levels of some long-chain acyl-carnitines, but not conjugated bile acids – still elevated in this subgroup of patients compared to the untreated group (**Supplementary Figure 2.G**).

### Comparison of SS patients to less clinically severe SC genotypes show trends consistent with beneficial effects of transfusion and hydroxyurea

We then performed a direct comparison of non-recently transfused SS patients to the metabolome of patients with less clinically severe genotype of SC (**Figure 2.A**). Linear models identified alterations of sphingolipid metabolism (especially sphingosine 1-phosphate, S1P and sphinganine 1-phosphate), methionine sulfoxide, phosphocreatine and PPP metabolites (gluconolactone phosphate and ribose diphosphate) as the top markers distinguishing SS and SC patients, even upon adjustment for HbA% (**Figure 2.B**). These findings were corroborated by HCA, which also showed enrichment in SS plasma of acyl-carnitines, taurine- and glycine-conjugated bile acids (taurodexycholate, taurocholate, taurochenodeoxycholate, taurolithocholate, glycocholate, glycochenodeoxycholate), carboxylic acids (succinate, malate, oxaloacetate), purine deamination products (urate, hydroxisourate), as opposed to depletion in L-arginine and S1P (**Figure 2.C**). These findings were further highlighted by the identification of metabolic correlates to HbC % (**Figure 2.D**), the top of which presented in the form of violin plots in **Figure 2.E**.

**Figure 2.**
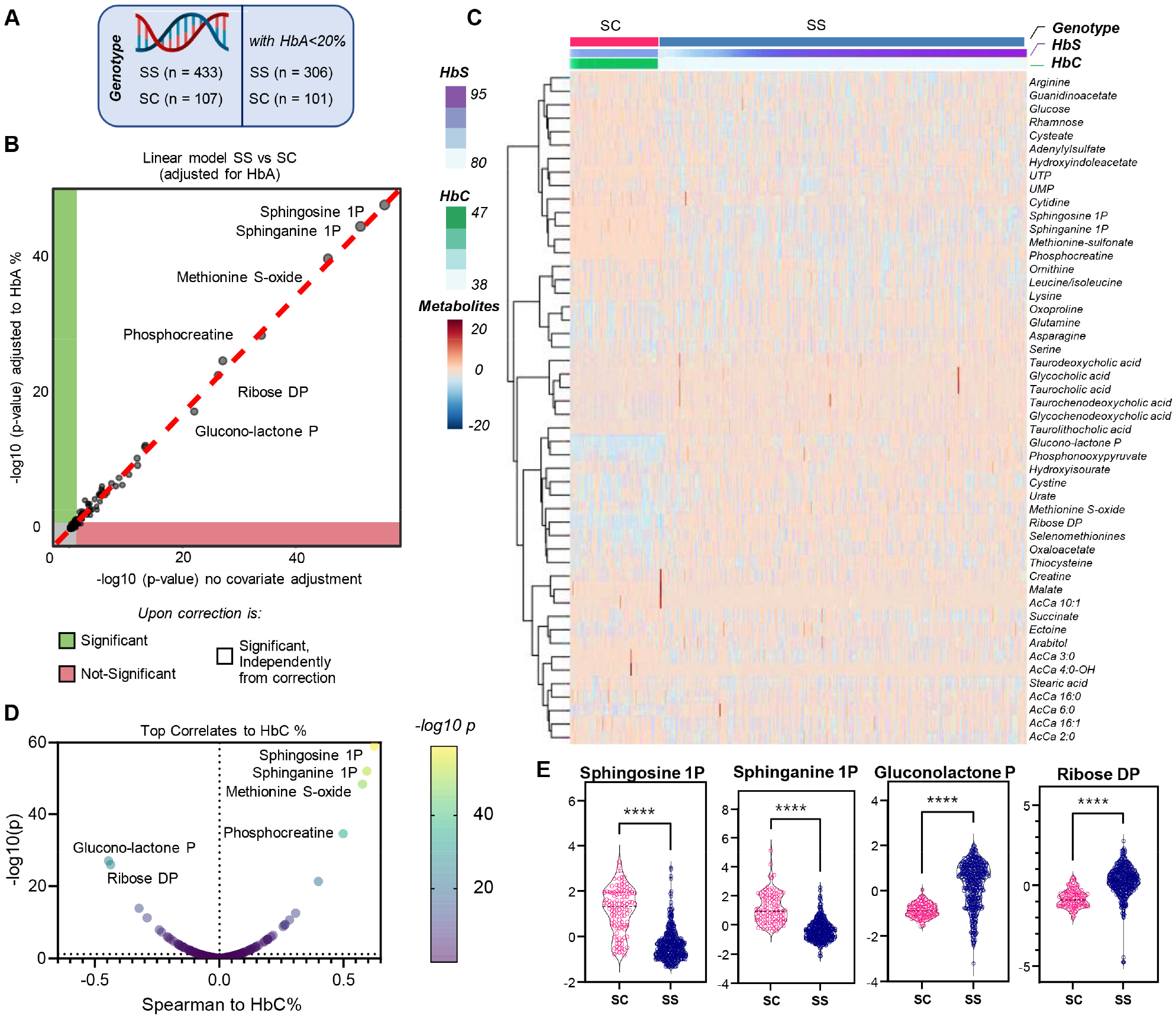
Plasma metabolic differences between SCD patients with SS and SC genotypes. Plasma metabolomics analyses were performed on plasma from patients with SS (n=433) or SC genotype (n =107 - **A**) who were not recently transfused (HbA <20% - n=306 and 101 for SS and SC, respectively). Linear models were used to identify the main differences between SS and SC patients, either unadjusted (x axis) or adjusted by HbA% (y axis - **B**). In **C**, hierarchical clustering analysis of the top 50 metabolites by t-test, and Spearman correlation analyses between plasma metabolites and HbC % (**D**). In **E**, focus on selected metabolites, including sphingosine 1-phosphate (S1P) and pentose phosphate pathway metabolites that differed significantly between plasma of SC and SS patients.

### Plasma metabolic markers of hemolysis in the SS SCD population

Degree of hemolysis was most elevated in SS patients with the more severe clinical manifestations. Analysis of metabolomics data as a function of the degree of hemolysis were thus performed in non-recently transfused (HbA<20%) SS patients via PCA, HCA and correlation analysis (**Figure 3.A-C**). Results indicate a strong positive correlation of hemolysis with circulating levels of conjugated bile acids, urate, acyl-carnitines, free fatty acids, pentose and hexose phosphate metabolites, carboxylic acids (succinate, malate) and a negative correlation with free amino acids, especially L-arginine and tryptophan metabolites (e.g., kynurenine). Similar results were even more evident in **Supplementary Figure 3**, in which we focused on the top 30 patients with highest and lowest overall hemolysis, without constraining for patients’ genotypes nor for recent transfusion events.

**Figure 3.**
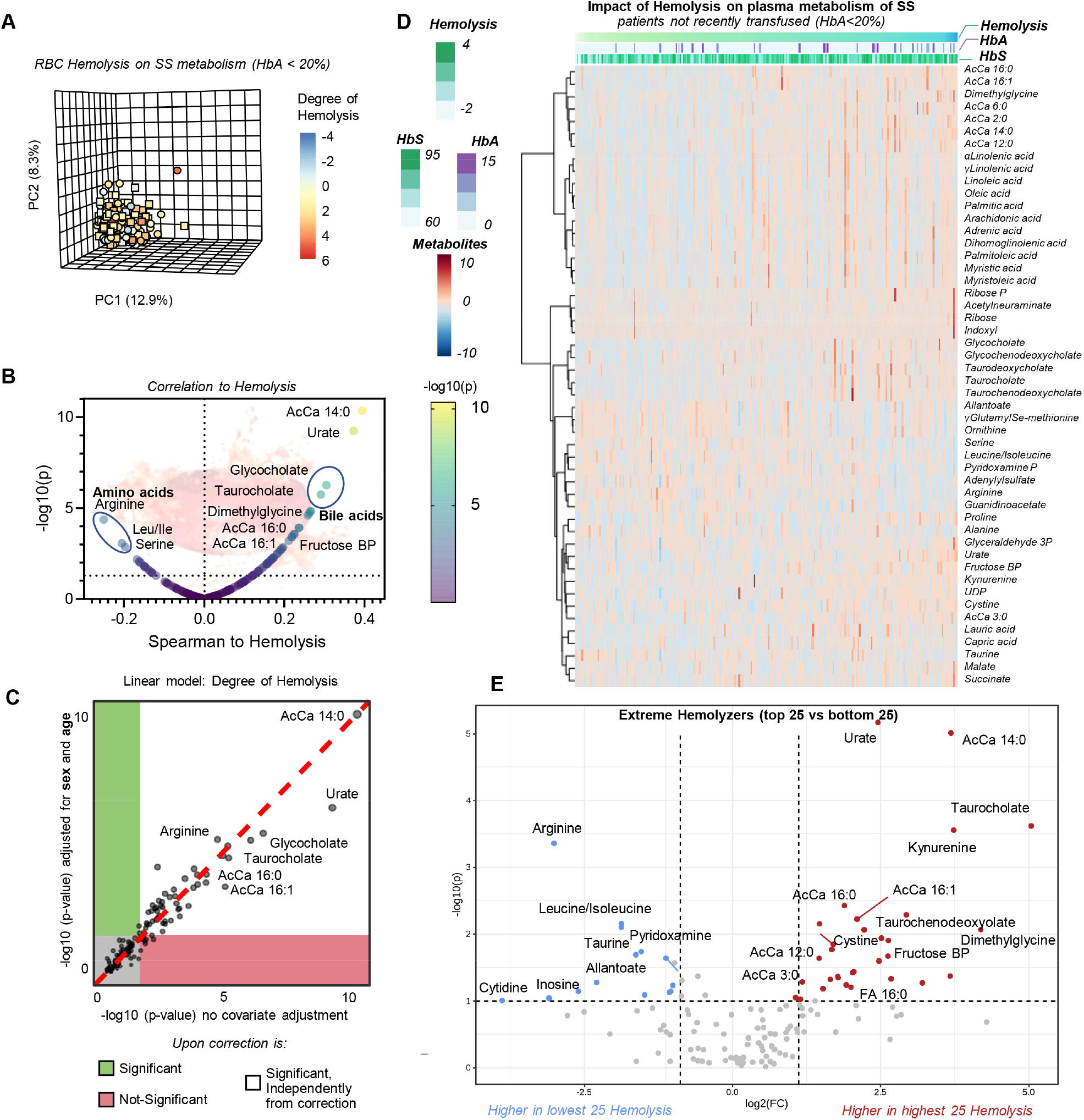
Plasma metabolic markers of hemolysis in SS SCD patients who were not recently transfused (HbA <20%) In **A**, principal component analysis of the plasma metabolomics data from this cohort, color-coded based on the degree of hemolysis. In **B, C** and **D** heat map, linear model (unadjusted or adjusted by subject sex and age on the y axis) and volcano plot of the plasma metabolites most significantly associated with hemolytic propensity. In **E**, volcano plot of the most significant plasma metabolic changes between the 25 patients with the highest and lowest degree of hemolysis (**F**).

### Plasma metabolic correlates to Doppler-echocardiographic tricuspid regurgitation jet velocity (TRV) in the WALK-PHaSST cohort

Cardiorenal dysfunction secondary to elevated hemolysis are common comorbidities in SCD, especially in patients with SS genotypes.^13,28^ Altered TRV is a hallmark of pulmonary dysfunction in this patient population, though plasma metabolic markers of aberrant TRV have not yet been reported, and is the focus of the analysis in **Figure 4**. Focusing on non-recently transfused (HbA<20%) SS patients, PCA, HCA, linear models (unadjusted, or adjusted for relevant covariates of patients’ age and sex) and correlation analyses all highlighted a significant correlation to TRV for plasma levels of S1P, L-arginine and tryptophan (depleted with high TRV) and their catabolites (citrulline, creatinine, kynurenine), acyl-carnitines (elevated with high TRV – **Figure 4.A-D**). Similar results were even more evident in **Supplementary Figure 4**, in which we focused on the top 30 patients with highest and lowest overall TRV, without constraining for patients’ genotypes nor for recent transfusion events. A snapshot of this analysis is provided in the volcano plot in **Figure 4.E**, which focuses on SS patients with high vs low TRV – further highlighting depletion of glutamine and 5-oxoproline (a product of the gamma-glutamyl-cycle) in patients with elevated TRV.

**Figure 4.**
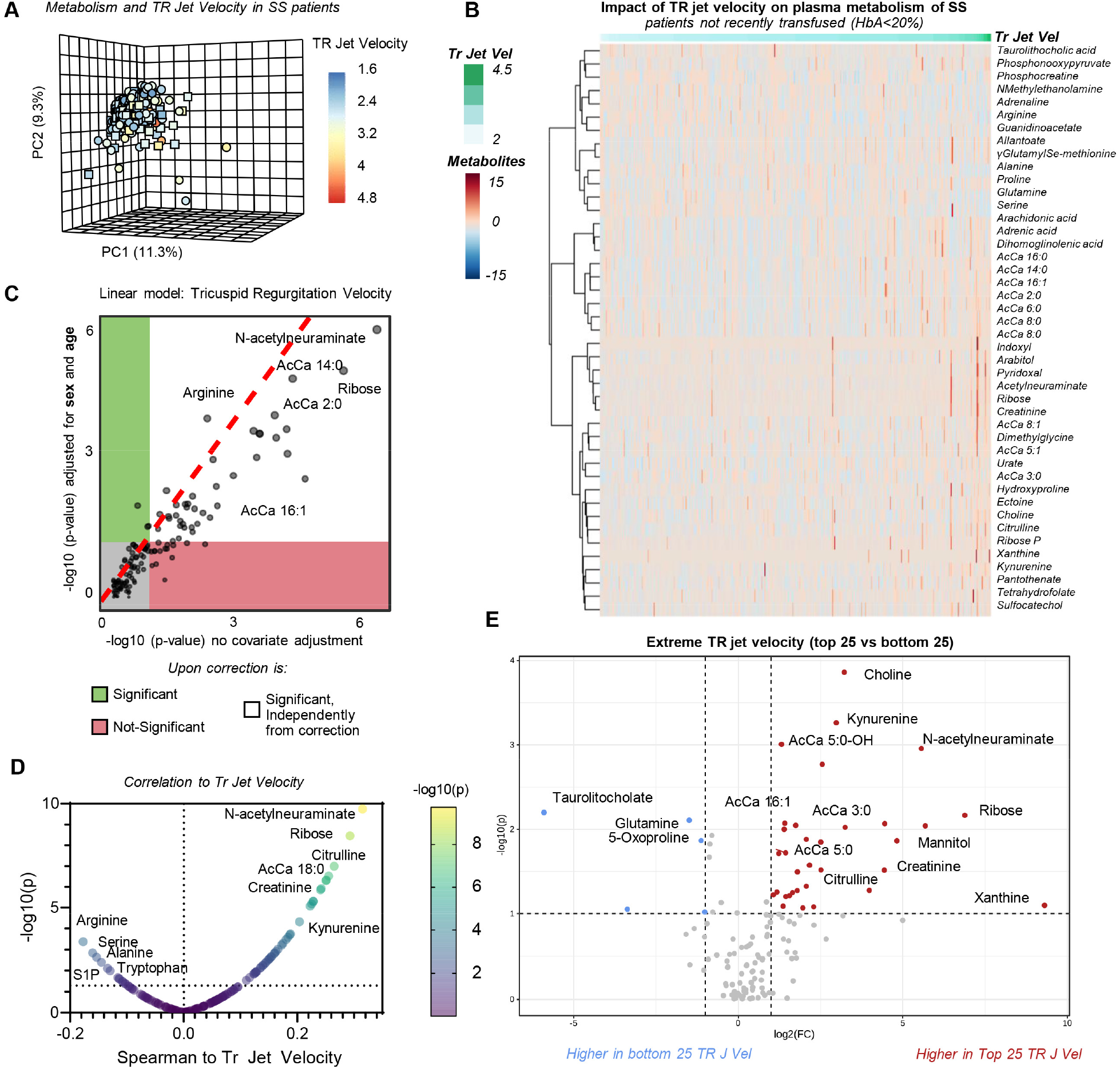
Plasma metabolic markers of cardiovascular dysfunction as gleaned by TR_jet velocity in SS patients. In **A**, principal component analysis of the plasma metabolomics data from SS patients with no recent transfusion event (HbA<20%), color-coded based on the tricuspid regurgitation velocity (TRV). In **B, C** and **D** heat map, linear model (unadjusted or adjusted by subject sex and age on the y axis) and volcano plot of the metabolites most significantlyassociated with TRV. In **E**, volcano plot and heat map of the most significant metabolic changes between the 25 patients with the highest and lowest degree of TRV.

### Plasma metabolic correlates to renal function in the WALK-PHaSST cohort

Plasma metabolic markers of renal function in SS patients who had not been transfused recently (HbA<20%) were here evaluated as a function of estimated glomerular filtration rates (eGFR), via PCA, HCA, linear models (unadjusted, or adjusted by patients’ sex and age) and correlation analyses (**Figure 5.A-D**). Results confirmed a significant negative correlation between eGFR and creatinine, with concomitant accumulation of acyl-carnitines, depletion of arginine and accumulation of its catabolites (creatinine, citrulline), elevation in the levels of pyridoxal, n-acetylneuraminate, pro-inflammatory carboxylic acids (succinate), kynurenine, choline, acetyl-choline, urate, indoles. Similar results were even more evident in **Supplementary Figure 5-6**, in which we focused on the top 25 patients with highest and lowest overall eGFR as a function of patients’ age in SS patients only with no recent transfusion event (HbA<20% - volcano plot in **Figure 5.E** and heat map in **Supplementary Figure 5**) or the top 30 patients by high vs low eGFR overall, without constraining for patients’ genotypes nor for recent transfusion events (**Supplementary Figure 6**).

**Figure 5.**
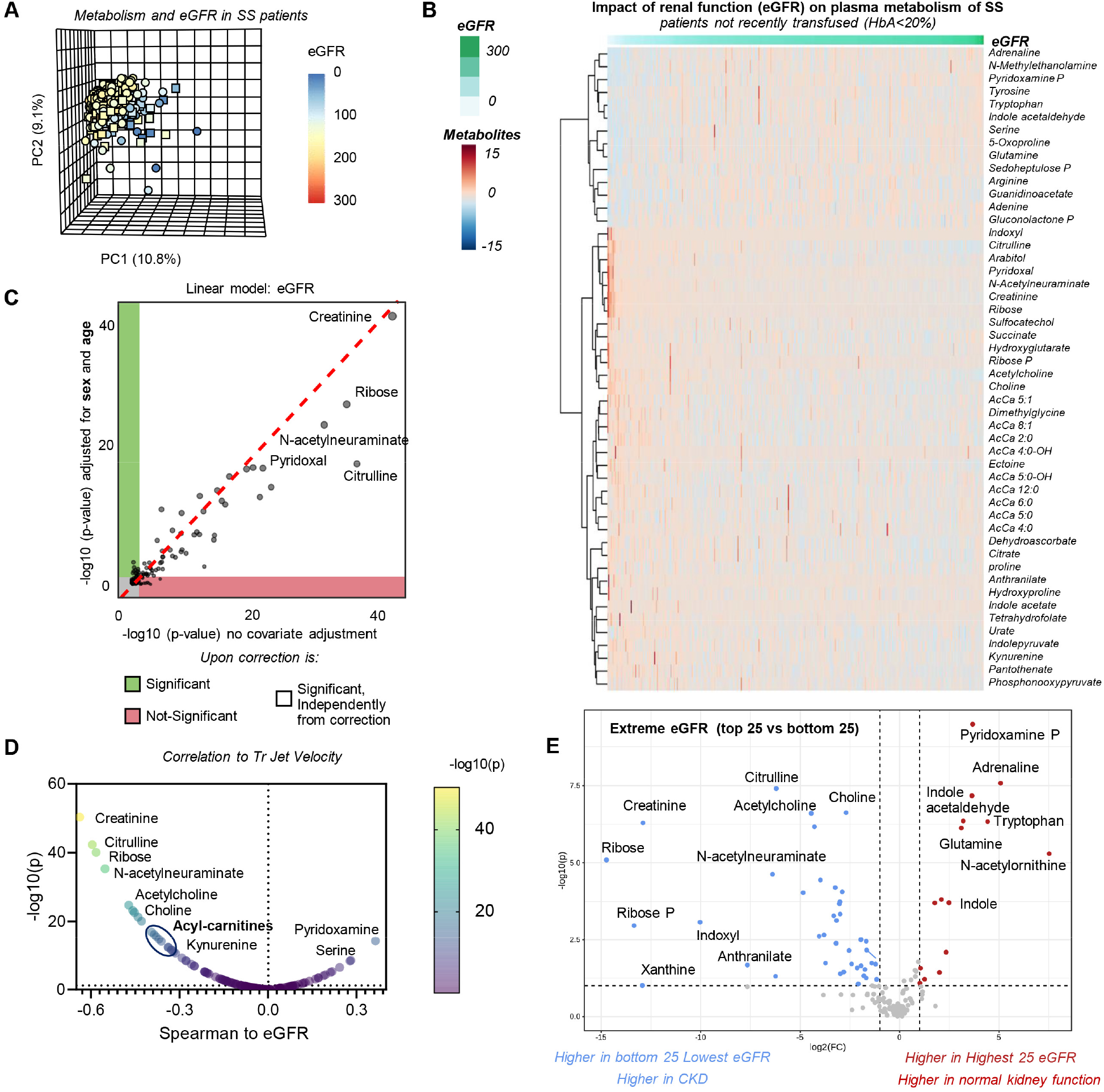
Plasma metabolic markers of renal dysfunction as gleaned by eGFR and creatinine in SS patients. In **A**, principal component analysis of the metabolomics data from this cohort, color-coded based on the eGFR. In **B, C** and **D**, heat map, linear model (unadjusted or adjusted by subject sex and age on the y axis) and volcano plot of the metabolites most significantly associated with eGFR. In **E**, volcano plot and heat map of the most significant metabolic changes between the 25 patients with the highest and lowest eGFR.

### Plasma metabolic markers of mortality, time to event and hazard ratio in the WALK-PHaSST cohort

Finally, we performed biomarker analyses to identify plasma metabolites associated with signatures of mortality, as gleaned by volcano plot of dichotomy outcomes (dead vs alive – **Figure 6.A**), receiver operating characteristics (ROC) curve determination (**Supplementary Figure 7**) and a more formal hazard ratio of metabolomics data adjusted by patients’ age (**Figure 6.B**). Results confirmed a strong association between conjugated bile acids, arginine and tryptophan metabolism (creatinine, citrulline, kynurenine, anthranilate, indoles), carboxylic acids (2-hydroxyglutarate, succinate), acyl-carnitines and untoward clinical outcomes. Correlation of clinical and metabolic covariates were thus plotted in the form of a network analysis in **Figure 6.C**, which identified hubs of conjugated bile acids, free fatty acids (especially very-long chain poly- and highly-unsaturated fatty acids), glutamine metabolites (glutamine, glutamate, 2-oxoglutarate), pantothenate and acyl-carnitines associated with hematological (HbS, HbA, HbC, HbF %; RBC, WBC counts) and cardiorenal function parameters (eGFR, creatinine, siten, BMI, cardiovascular risk, hemolysis). Further correlation analyses are reported for these clinical covariates to plasma metabolite levels in **Supplementary Figure 8**.

**Figure 6.**
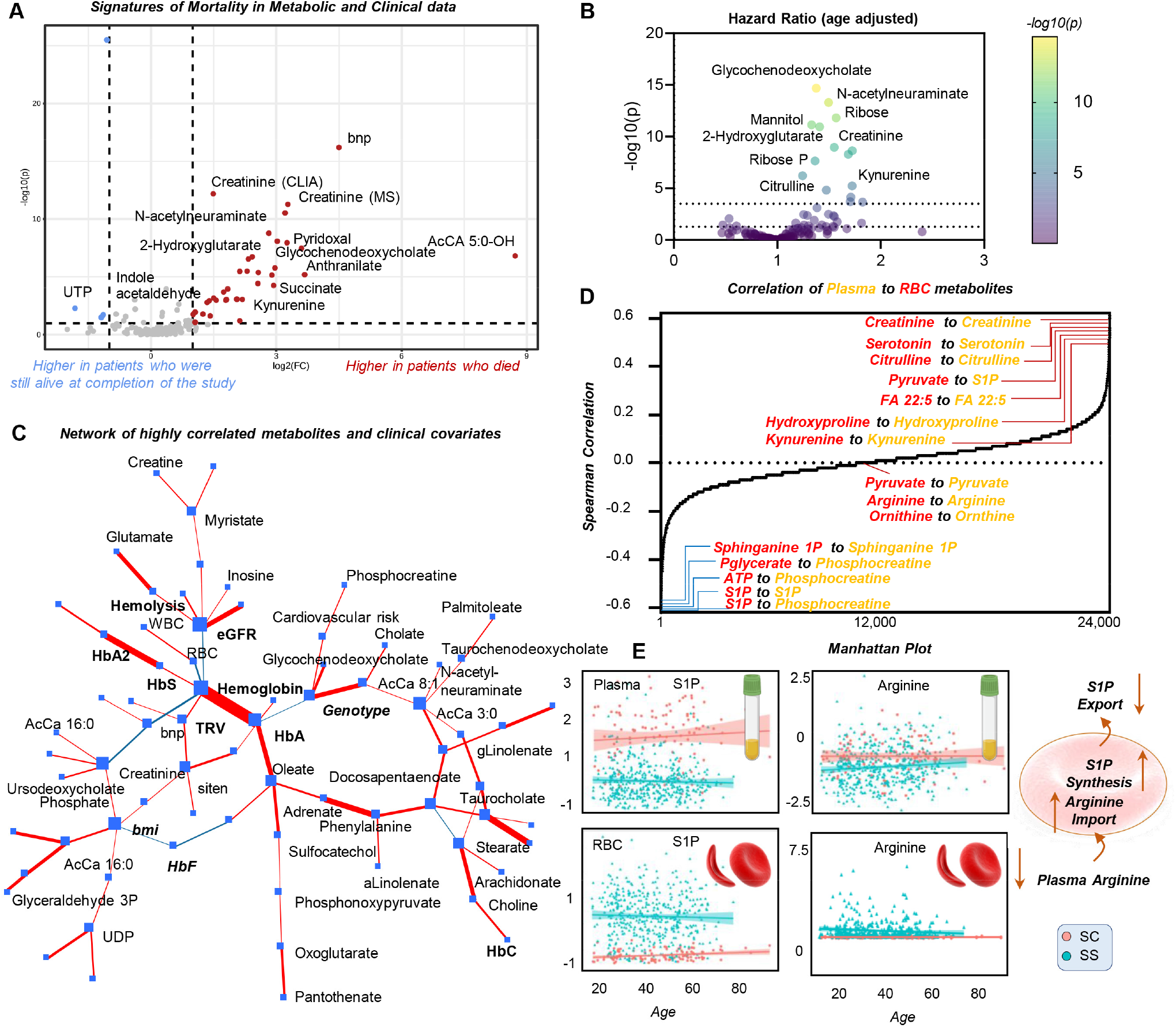
Plasma metabolic signatures of mortality, hazard ratio and plasma metabolic correlates to previously published RBC metabolomics data in the WALK-PHaSST cohort. Plasma metabolic correlates to mortality in the WALK-PHaSST cohort (**A**). In **B**, results from the hazard ratio analysis of plasma metabolomics data adjusted by patients’ age. In **C**, plasma metabolite levels from the present analysis of 596 samples were correlated with all available clinical and hematological covariates and plotted in the form of a network, where each node represents a covariate, and edges indicate correlation between two covariates (thickness proportional to the module and significance of this association). In **D**, plasma metabolite levels from the WALK-PHaSST cohort were matched to RBC metabolomics measurements (*D’Alessandro et al*.^*23*^). Results indicated a strong positive cross-matrix correlation for almost all metabolic markers of cardiorenal dysfunction, except for sphingosine 1-phosphate (S1P), whose levels were elevated in SS RBCs and SC plasma, suggesting an increased synthesis or decreased export in the more clinically severe SS genotype (**E**).

### The Sickle Cell Disease online portal for real time processing of plasma and RBC metabolomics data in SCD against clinical covariates, and cross-matrix correlations

Clinical and plasma metabolomics data in this study, and RBC metabolomics data from our previous study^23^ were collated into an online portal for real time data processing and figure generation. The portal, open and accessible at https://mirages.shinyapps.io/SCD2023/ serves as a repository for the data generated as part of this study, and as a tool to perform real time analyses and export of results upon correlation analyses between metabolites and clinical covariates, with the opportunity to modify filters (e.g. age, sex) or cross-matrix (plasma vs RBC) correlations. The portal offers the opportunity to correct for covariates of interest, for example by adjusting by HbS, HbA, HbF and HbC% (**Supplementary Figure 9**). Leveraging this portal we performed cross-matrix analysis of plasma and RBC metabolomics data from the Walk-PHASST study. An overview of the matrix of plasma and RBC intra- and cross-matrix metabolic correlates is provided in **Figure 6.D**, where metabolite pairs (x axis) are sorted by Spearman correlation coefficients (from lowest to highest). These analyses are further presented in the correlation matrix in **Supplementary Figure 10**. Notably, most metabolic markers of cardiorenal dysfunction (especially creatinine, citrulline and tryptophan metabolites, serotonin and kynurenine) showed significant (p<0.0001) positive correlations (r>0.5) between plasma and RBC levels. Arginine and ornithine levels, like pyruvate, poorly correlated between plasma and RBCs, suggesting cell-intrinsic mechanisms affecting the levels of these metabolites in SCD. However, these analyses also identified a discrepancy between elevation of intracellular RBC S1P levels and depletion of circulating S1P in plasma of patients with SS genotype compared to any other genotype tested in this study (**Figure 6.E; Supplementary Figure 11**). Similarly, increased arginine uptake from plasma to RBCs could explain the observed compartment and genotype-specific (SS vs SC) changes in arginine levels (**Figure 6.E; Supplementary Figure 11**).

## DISCUSSION

Here we performed an extensive plasma metabolomics analysis of a clinically well-curated cohort of 596 SCD patients. Availability of detailed clinical measurements of HbS, A, C and F percentages allowed us to identify the impact of recent transfusion events, genotype (with an emphasis on the more severe SS vs the less clinically severe SC genotype). As a result, we identified multiple signatures associated with cardiorenal dysfunction and hazard ratios, including alterations in tryptophan/kynurenine/indole metabolism, bile acid deconjugation, acyl-carnitine and S1P metabolism, glutaminolysis and mitochondrial dysfunction and L-arginine metabolism to citrulline and creatinine (**Figure 7**). Results are relevant in that they confirm and expand upon previous findings in the literature, linking metabolic differences^23^ to clinically relevant outcomes and providing a metabolic window on clinical efficacy of common interventions like transfusion of packed RBCs and treatment with hydroxyurea.^39^ For example, previous smaller scale studies had documented a beneficial effect of transfusion events in SCD in the circulating plasma and RBC metabolome,^40^ though the scale of these studies was limited to a handful of recipients with no characterization of clinical relevant covariates.

**Figure 7.**
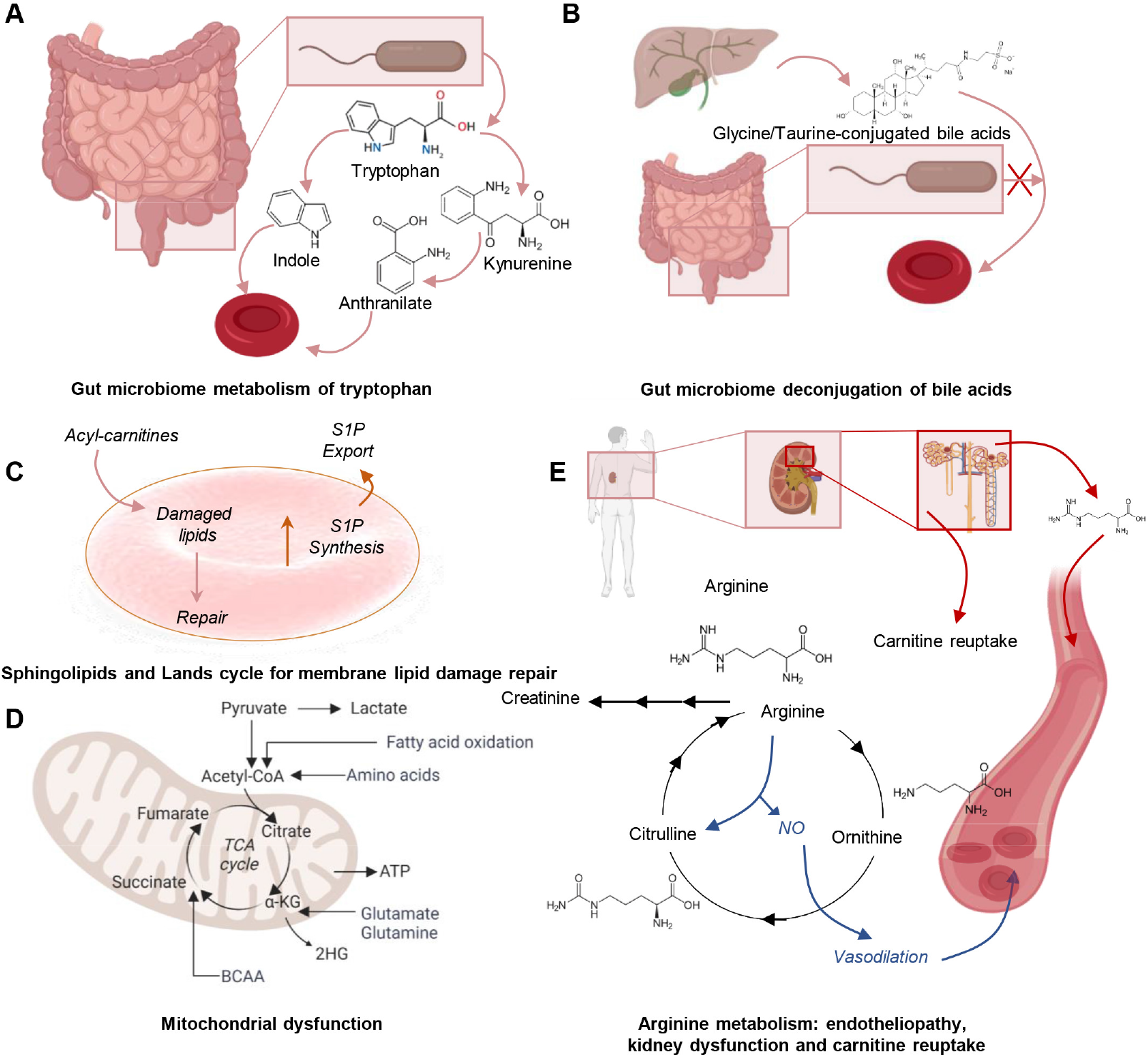
Summary of the main metabolic pathways affected in plasma of SCD patients that correlate with relevant clinical outcomes. Plasma metabolomics analyses identified dysregulation of tryptophan/kynurenine/indole metabolism (**A**), conjugated bile acids (**B**), elevation of acyl-carnitines (**C**), mitochondrial dysfunction (**D**), arginine metabolism (**E**) as critically altered metabolic pathways associated with cardiorenal dysfunction and hazard ratios in this patient population.

Previous studies had identified dysregulated arginine metabolism^41^ via hemolysis-triggered increases in arginase activity as factor limiting nitric oxide synthesis and vasodilation in SCD patients, ultimately contributing to endothelial and cardiovascular dysfunction.^13^ Here we expand on these findings identifying a role for transfusion events or hydroxyurea treatment in the amelioration of dysregulated metabolic signatures in these pathways, effects that result in the mitigation of arginine consumption and accumulation of creatinine, ornithine, citrulline and guanidinoacetate.

Similarly, previous studies had focused on glutamine consumption in the context of SCD, and glutamine supplementation as a therapeutic strategy to fuel de novo glutathione synthesis as a strategy to counteract oxidant stress in the sickle RBC, though findings are controversial as to whether glutamine supplementation aggravates or mitigates the incidence of pain and vasocclusive crises in this patient population.^17,18,42^ Our findings show that accumulation of carboxylic acids – especially succinate and malate – are observed in plasma concomitantly to glutamine consumption and generation of 5-oxoproline, a byproduct of glutathione catabolism via the gamma-glutamyl-cycle and a metabolic end-product of this cycle in the mature RBC, owing to the lack of the enzyme oxoprolinase. This observation is relevant in light of the role of these carboxylic acids, especially succinate, in the stabilization of hypoxia inducible factor 1 alpha,^43^ and activation of its transcriptional targets, including interleukin 1beta.^44^ Of note, accumulation of amino and carboxylic acids downstream to glutamine catabolism (glutamate, 2-oxoglutarate, 2-hydroxyglutarate, succinate – in particular) was linked to signatures of cardiorenal dysfunction and overall hazard ratio, suggesting that fueling of glutathione synthesis via oral supplementation not only promotes anaplerotic reactions, but also likely fuels catabolism towards the accumulation of pro-inflammatory small molecule mediators. Further studies with stable isotope-labeled tracers will be necessary to further test this hypothesis and the ensuing clinical ramifications.

Glycine and taurine conjugated bile acids are synthesized in the liver and deconjugated by the gut microbiome, and their accumulation in plasma is consistent with microbiome dysbiosis.^45,46^ These metabolites have been implicated in the pro-inflammatory activation of macrophages,^47^ as well as in coagulopathic complications (e.g., after trauma and hemorrhage),^48^ both common comorbidities in sickle cell patients. Conjugated bile acids had been previously identified as markers of cholecystectomy in sickle cell patients,^42^ though the impact of patients’ genotypes, transfusion events or hydroxyurea treatment on the circulating levels of these metabolites had not been reported. Of note, indole metabolites derived from tryptophan catabolism are also derived from microbial catabolism^49^, and participate in systems homeostasis.^50^ Like kynurenines,^51^ other endogenous tryptophan metabolites, indoles participate in immunomodulatory effects on macrophages through triggering of aryl hydrocarbon receptors.^52^ Of note, kynurenine was here associated with high hemolysis, increased TRV and hazard ratios. This finding is interesting in that elevated kynurenine was previously reported in individuals suffering with elevated cardiopulmonary dysfunction, pulmonary hypertension and lung fibrosis, such as subjects with Down syndrome.^53^ Elevation of circulating levels of kynurenine is a hallmark of viral infection with pulmonary complications, such as COVID-19, whereby this metabolite correlates with interleukin 6 levels, clinical severity of SARS-CoV-2 infection and, ultimately, clinical outcomes.^54,55^ Of note, in both cases elevation of kynurenine is associated with the activation of the interferon-indoleamine 2,3-dioxygenase axis, in the latter case stimulated by sensing of viral nucleosides by the cGAS-STING system.^56^ It is interesting to note that mitochondrial DNA and RNA (e.g., in the context of aging) have been reported to contribute to activation of the cGAS-STING responses.^57^ Since mature sickle RBCs have been reported to contain up to 5-6 residual mitochondria per cell,^58,59^ in light of the elevated degree of sickle RBC hemolysis, it is tempting to speculate that the elevated kynurenine levels as a function of high degree of hemolysis reported here participate in a similar circulating mitochondrial DNA sensing cascade, which could pave the way for potential novel therapeutic interventions. Finally, it is worth noting that elevation of kynurenine and indoles subtracts a substrate for the synthesis of serotonin, which is an essential component of platelet dense granules^60^ and a neurotransmitter that could contribute to counteracting nociceptive signaling.^61^

Here we performed network analyses of combined metabolomics and clinical data, which showed a strong association between hematological (e.g., HbS%) and clinical parameters with all metabolites mentioned in this discussion, as well as free fatty acids and acyl-carnitines. Of note, elevated kynurenine levels had been previously associated with increased susceptibility to hemolysis in the context of obesity,^62^ whereby depletion in circulating free fatty acids (previously also reported in pediatric patients with SCD^63^) and elevation acyl-carnitines had been identified as a marker of increased membrane RBC lipid damage and remodeling. This observation was in keeping with previous reports on the activation of the so-called Lands cycle in the context of SCD^64^ (or even just sickle cell trait)^65^, or other osmotic, mechanical or oxidative stressors to RBCs, such as blood storage,^66^ exercise,^67^ chronic^68^ or acute kidney dysfunction.^69^ Of note, our findings expand on these observations, by identifying acyl-carnitine accumulation (and correction thereof by transfusion more than hydroxyurea treatment) as a hallmark of cardiorenal dysfunction in the sickle cell patient population. Interventions with carnitine supplementation in the context of SCD^70^ may thus be beneficial – especially in combination with hydroxyurea^71^ - to fuel the Lands cycle and contribute to prevent or mitigate RBC membrane lipid damage.

Here we report a significant association between S1P depletion in plasma and genotypes with distinct severity of clinical manifestations (SS vs SC). In previous studies, we and others had reported an elevation of RBC levels of the sphingolipid S1P in association with disease severity^22^ or increased (normal and sickle) erythrocyte susceptibility to extravascular hemolysis.^72^ While the mechanism of action is unclear, it has been posited that S1P may contribute to 2,3-diphosphoglycerate-dependent modulation of oxygen off-loading.^22^ By contributing to deoxyhemoglobin stabilization, hypoxia-induced elevation of S1P^73^ may cooperate with DPG to impact the so-called oxygen-dependent metabolic modulation that involves glycolytic enzyme sequestration to the membrane via binding to band 3, which in turn regulates RBC capacity to activate the PPP to generate reducing equivalents to sustain antioxidant challenges.^74,75^ However, here we observe a significant depletion of circulating S1P in plasma from subjects with SS genotypes, especially upon correction for transfusion events or hydroxyurea treatment. These results are suggestive of a role for decreased export, rather than (or in combination with) elevated synthesis may be responsible for dysregulated S1P metabolism in SCD. These results could be perhaps explained by polymorphisms in the S1P transporter Mfsd2b^76^ – previously associated with elevated susceptibility to osmotic fragility in healthy blood donor volunteers and sickle cell patients.^37^

In brief, our study expands on existing literature and provides an extensive overview of the plasma metabolome in SCD and its clinical correlates, to the extent they are affected by common interventions like blood transfusion or treatment with hydroxyurea. The generation of a novel online portal as a repository for all the data described in this study will facilitate further dissemination and elaborations of the data presented herein in the future.

## Acknowledgments

AD was supported by funds by the National Institute of General and Medical Sciences (RM1GM131968), and from the National Heart, Lung, and Blood Institute (R01HL146442, R01HL149714, R01HL148151, R01HL161004). MTG receives research support from NIH grants 5R01HL098032, 2R01HL125886, 5P01HL103455, 5T32HL110849, and UH3HL143192. This work was supported by UH3HL143192. The content is solely the responsibility of the authors and does not necessarily represent the official views of the National Institutes of Health.

## Authors’ contributions

SMN, YZ, VG, MTG designed and executed the WALK-PHASST study, collected and stored the samples, performed clinical measurements. FC, FG, JAR, AD performed metabolomics analyses (untargeted and targeted quantitative). KB, MG and JME built the SCD metabolome portal. AD performed data analysis and prepared figures and tables. AD wrote the first draft of the manuscript, which was revised by all the other authors. All the authors contributed to finalizing the manuscript.

