## Supplementary Methods and Figures for "Metabolic signatures of cardiorenal dysfunction in plasma from sickle cell patients, as a function of therapeutic transfusion and hydroxyurea treatment"

**SUPPLEMENTARY DATA TABLE** *(PLEAE REFER TO "LEGEND" SHEET WITHIN THE FILE)* .......... XLSX

### *Supplementary Materials and Methods - Extended*

***Ultra-High-Pressure Liquid Chromatography-Mass Spectrometry (MS) metabolomics:*** Frozen plasma aliquots (50 µL) were thawed on ice then extracted 1:25, in ice cold extraction solution (methanol:acetonitrile:water 5:3:2 *v/v/v*). Samples were vortexed for 30 min at 4℃ and insoluble material pelleted via centrifugation at 15,000 *g* for 15 min under refrigerated conditions, as described. [1,2] Analyses were performed using a Vanquish UHPLC coupled online to a Q Exactive mass spectrometer (Thermo Fisher, Bremen, Germany). Samples were resolved as described,[3,4] over a Kinetex C18 column (2.1x150 mm, 1.7 µm; Phenomenex, Torrance, CA, USA) at 45°C. A volume of 10 ul of sample extracts was injected into the UHPLC-MS. Each sample was injected with two different chromatographic and MS conditions as follows: 1) using a 5 minute gradient at 450 µL/minute from 5-95% B (A: water/0.1% formic acid; B:acetonitrile/0.1% formic acid) and the MS was operated in positive mode and 2) using a 5 minute gradient at 450 µL/minute from 5-95% B (A: 5% acetonitrile, 95%water/1 mM ammonium acetate; B:95%acetonitrile/5% water, 1 mM ammonium acetate) and the MS was operated in negative ion mode. The UHPLC system was coupled online with a Q Exactive scanning in Full MS mode at 70,000 resolution in the 60-900 m/z range, 4 kV spray voltage, 15 sheath gas and 5 auxiliary gas. These chromatographic and MS conditions were applied for both relative and targeted quantitative metabolomics measurements, with the differences that for the latter targeted quantitative post hoc analyses were performed on the basis of the stable isotope-labeled internal standards used as a reference quantitative measurement, as detailed below.

***Quality control and data processing:*** Calibration was performed prior to analysis using the Pierce™ Positive and Negative Ion Calibration Solutions (Thermo Fisher Scientific). Acquired data was then converted from raw to .mzXML file format using RawConverter. Samples were analyzed in randomized order with a technical mixture injected every 15 samples to qualify instrument performance and ensure technical coefficients of variations (standard deviation divided by the mean) below 20%. Metabolite assignments, isotopologue distributions, and quantification of stable isotope-labeled internal standards were performed using MAVEN (Princeton, NJ, USA), as described.[5]

***Statistical analyses:*** Data analysis was performed through the auxilium of the software MAVEN. Graphs and statistical analyses (either two-way ANOVA or repeated measures

ANOVA) were prepared with GraphPad Prism 9.0 (GraphPad Software, Inc, La Jolla, CA), GENE E (Broad Institute, Cambridge, MA, USA), Multivariate analyses, including principal component analyses (PCA), hierarchical clustering analyses, two-way ANOVAs, correlation analyses (Spearman) and calculation of receiver operating characteristic (ROC) curves were performed through the software MetaboAnalyst 5.0.[6] For survival analysis, we used time to right censorship (including death or last follow-up) as time to event and vital status (death or alive) was the studied outcome. PCA was used to derive a hemolytic component from lactate dehydrogenase, aspartate aminotransferase total bilirubin and reticulocyte percent.[7-9] We applied Cox proportional hazard models to calculate the hazard ratios and P value for each metabolite. and Cox analysis and adjusted regressions of metabolites to clinical outcomes were performed in R (R Core Team (2022), https://www.r-project.org/).

***Sickle Cell Disease ShinyApp portal*** A portal for online sharing of all the data generated in this study, in like-wise fashion to our recent COVID-Ome Explorer portal.[10] After data curation and quality control, each of the datasets (plasma metabolomics, RBC metabolomics from our previous study on the WALK-PHASST Cohort[11], clinical data) was linked at the sample level with a unique identifier, enabling cross-referencing among datasets. Then, each of the datasets was imported into applications developed using R, R Studio, and the R-based web application framework Shiny. All code required to run the COVIDome Explorer applications can be found at https://github.com/cusom/CUSOM.COVIDome.Shiny-Apps (Zenodo https://doi.org/10.5281/zenodo.5081091) and https://github.com/cusom/CUSOM.ShinyHelpers (Zenodo https://doi.org/10.5281/zenodo.5081093).

### SUPPLEMENTARY FIGURES

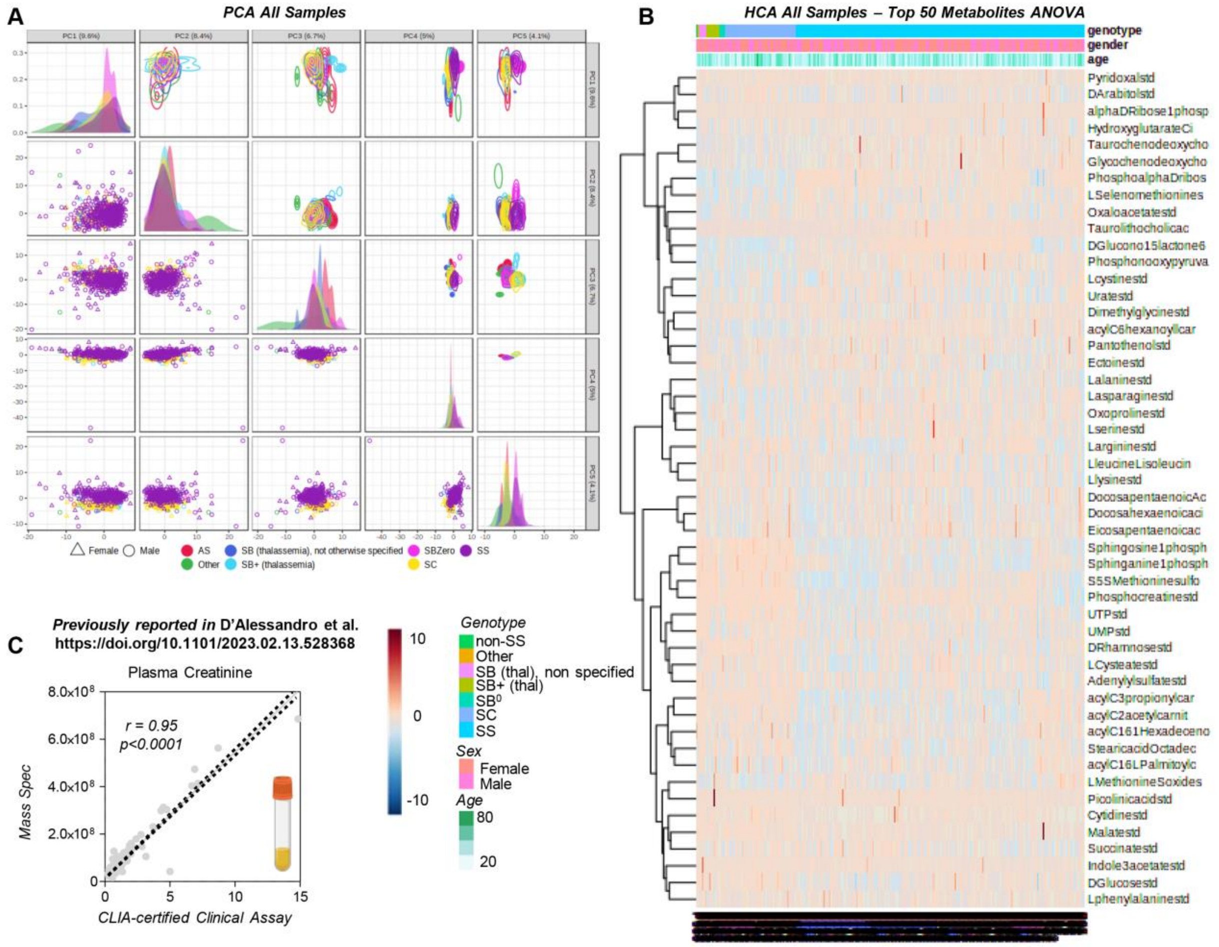

**Supplementary Figure 1** ***Plasma metabolomics of the WALK-PHASST SCD cohort*** Principal component analysis (PCA) of metabolomics data on 596 plasma samples (**A**), from 6 different genotypes (**B**). Of note, metabolomics measurements of plasma creatinine showed significant positive correlation (r=0.95) with CLIA-measurements in the clinics, despite double randomization of the samples, blinding of the mass spec lab and prolonged storage of the Walk-PHASST samples (**C**), as previously noted.[11]

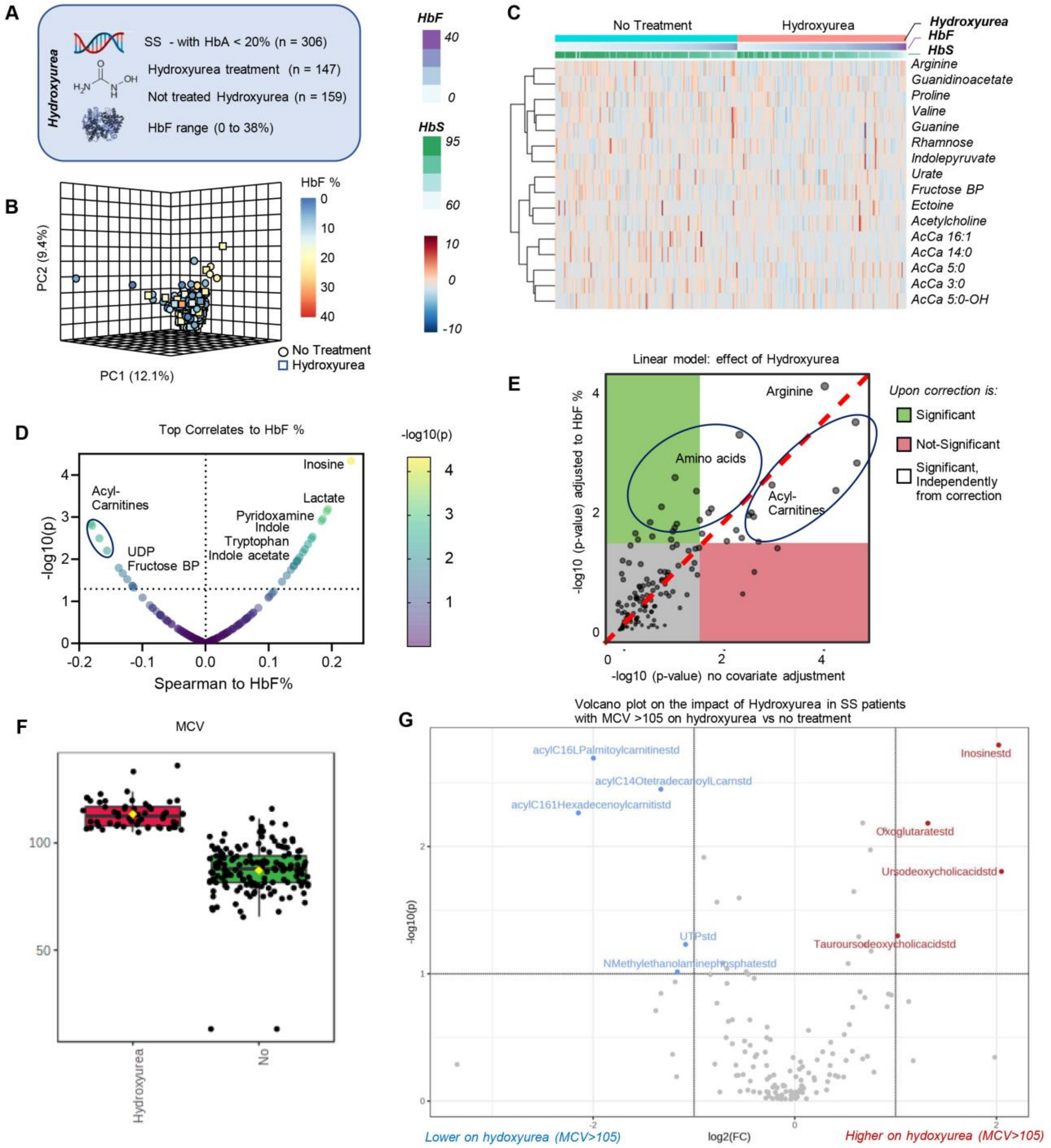

**Supplementary Figure 2** ***Plasma metabolic correlates to Hydroxyurea treatment in the WALK-PHaSST cohort*** SS patients who were not recently transfused (HbA<20%; n=306) were tested as a function of treatment with hydroxyurea (n=147 vs n=159 untreated - **A**). In **B**, principal component analysis of the data from this cohort, color-coded based on HbF %. In **C**, heat map of the most significant metabolites affected by the treatment. In **D**, metabolic correlates to HbF%. In **E**, linear model identifying the effect of hydroxyurea in SS patients, either unadjusted (x axis) or adjusted by HbF% (y axis). In **F**, to reduce the noise in the hydroxyurea group, owing to questionable adherence to the treatment, we narrowed the analysis down to the patients on treatment with MCV >105. The analysis (**G**) showed a beneficial effect of hydroxyrea in lowering the levels of several circulating acylcarnitines, an effect that was masked by the general analysis.

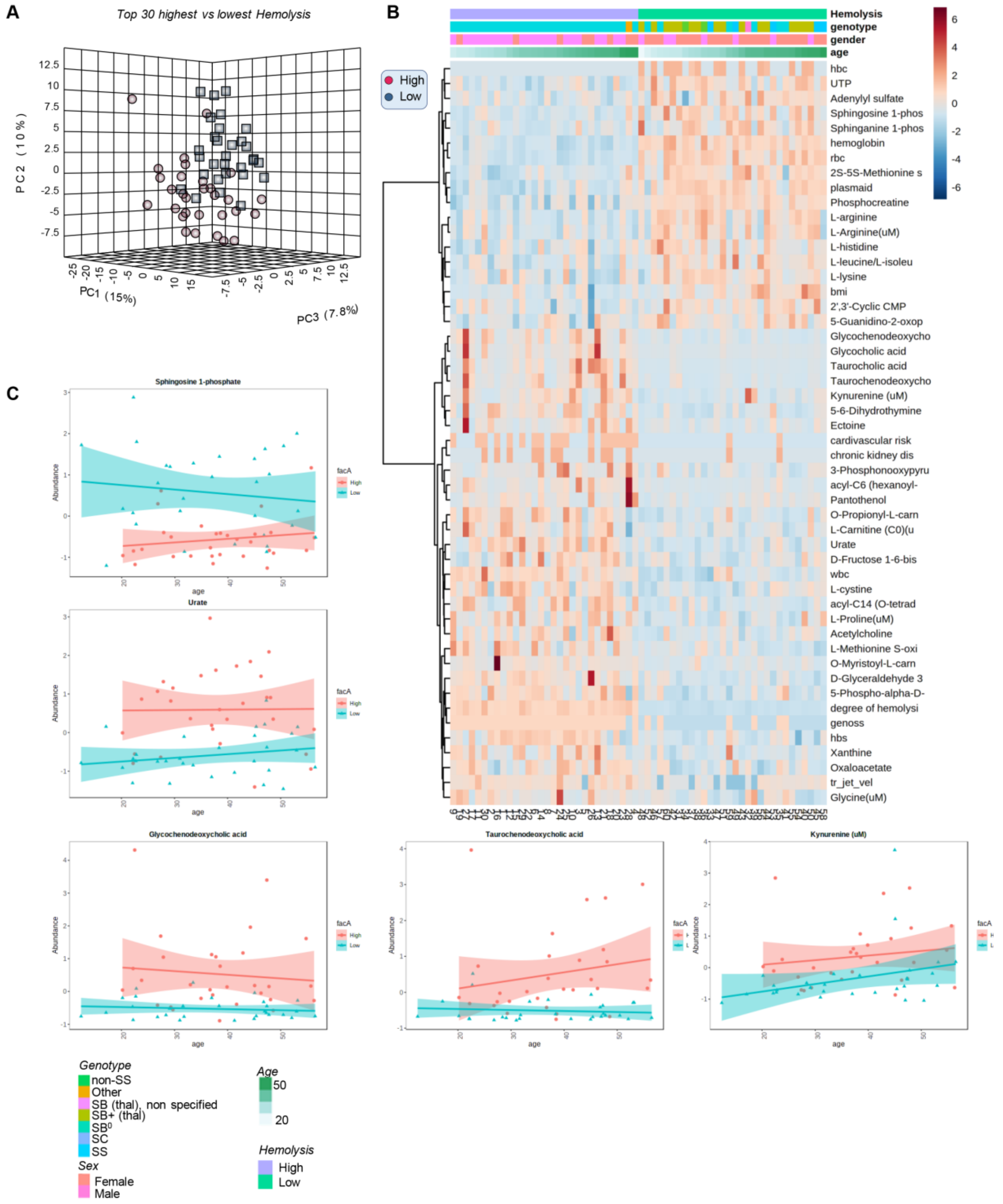

**Supplementary Figure 3** ***Plasma metabolomics of patients enrolled in the WALK-PHASST study with high and low hemolytic propensity*** tracks with genotypes (high in SS, low in SC) and shows distinct metabolic phenotypes, as gleaned by principal component analysis (**A**), hierarchical clustering analysis (**B**) and line plots (**C** – x axis tracks with patients' age).

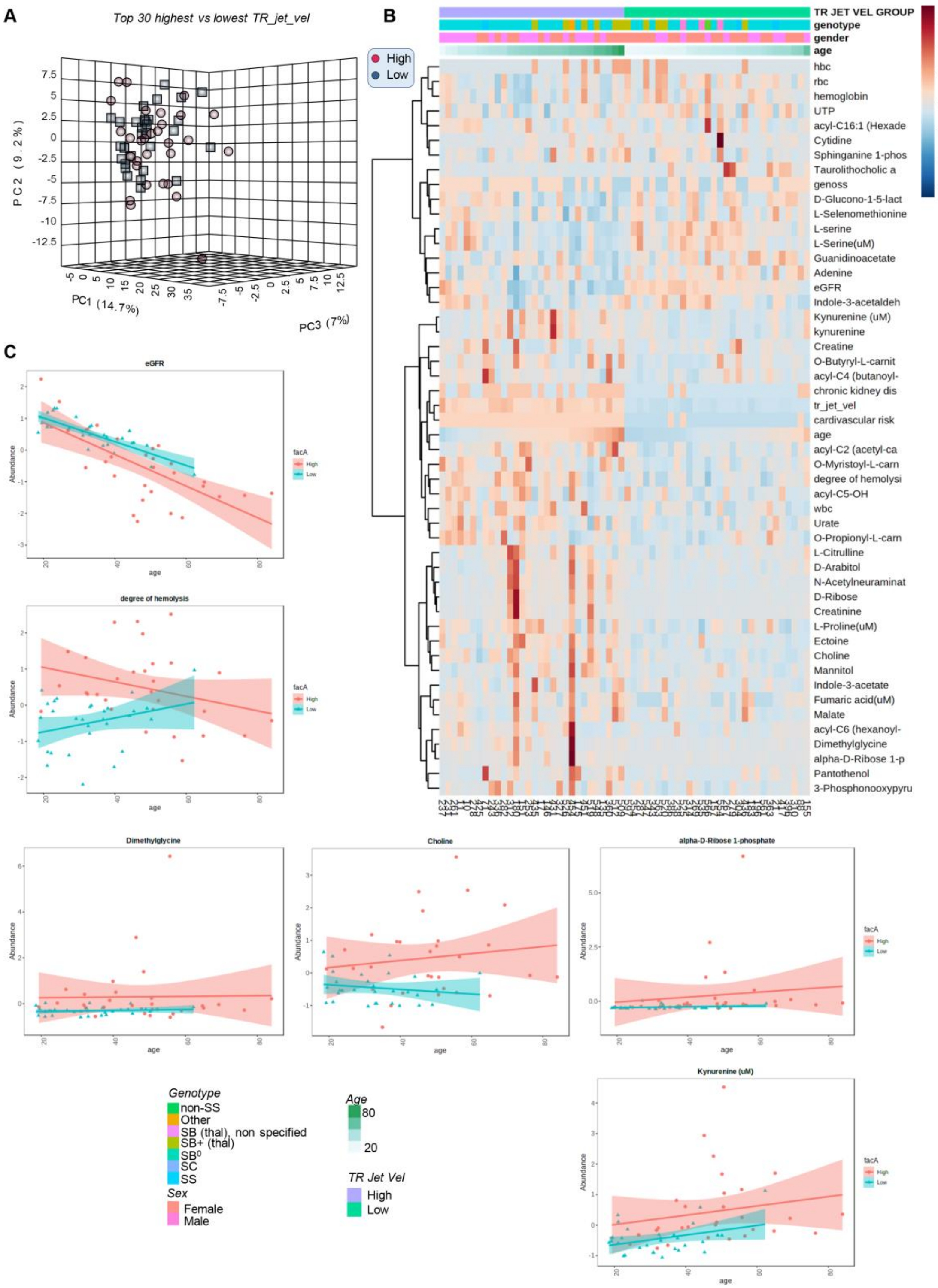

**Supplementary Figure 4** ***Plasma metabolomics of patients enrolled in the WALK-PHASST study with high and low tricuspid regurgitation velocity (TRV)*** tracks with genotypes (high in SS, low in SC), age (high in older patients) and shows distinct metabolic phenotypes, as gleaned by principal component analysis (**A**), hierarchical clustering analysis (**B**) and line plots (**C** – x axis tracks with patients' age).

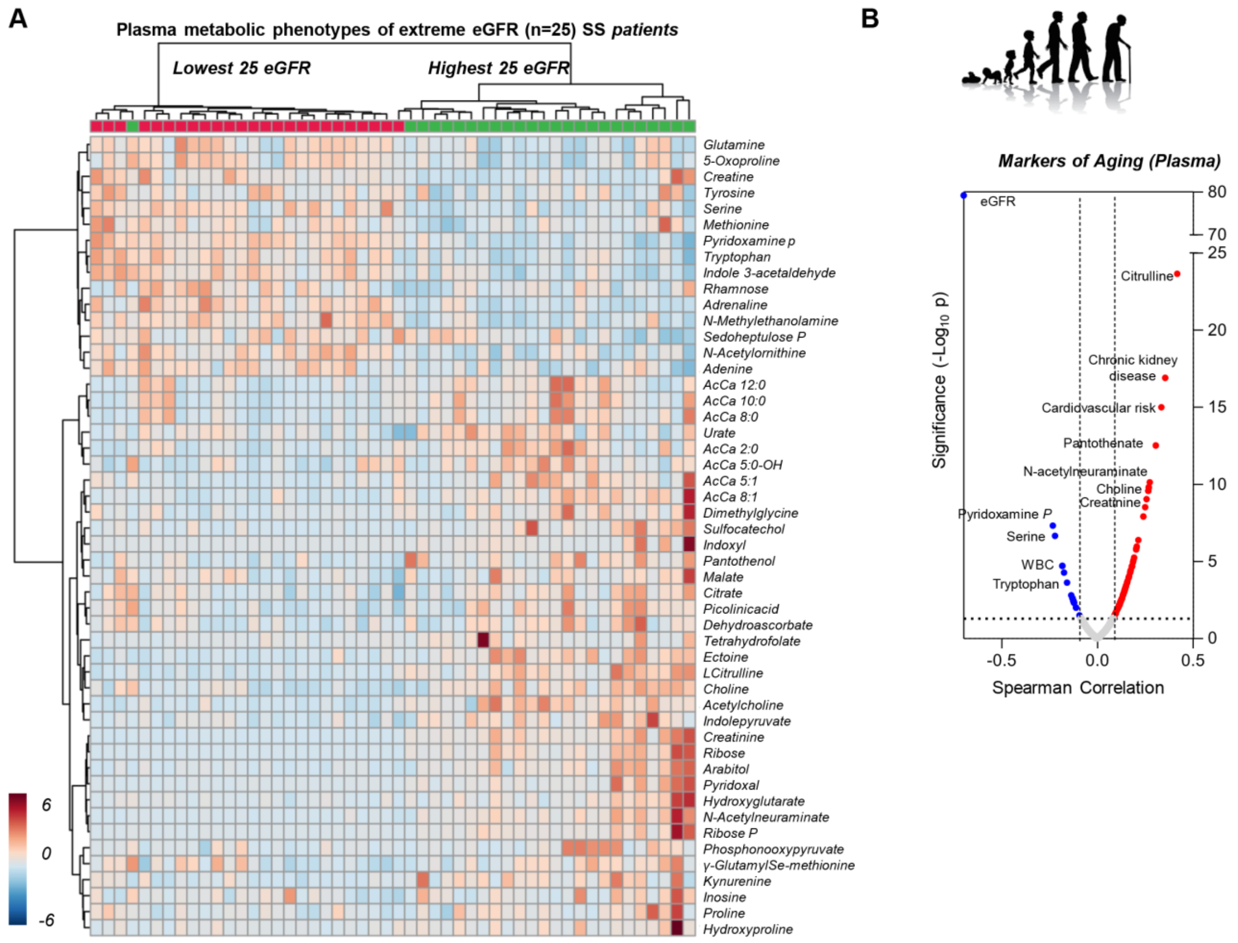

**Supplementary Figure 5** ***Plasma metabolomics of patients enrolled in the WALK-PHASST study with high and low eGFR propensity***, as gleaned by hierarchical clustering analysis (**A**). Metabolic markers and clinical measurements of kidney dysfunction were identified as the top correlates to patients' age (Spearman - **B**).

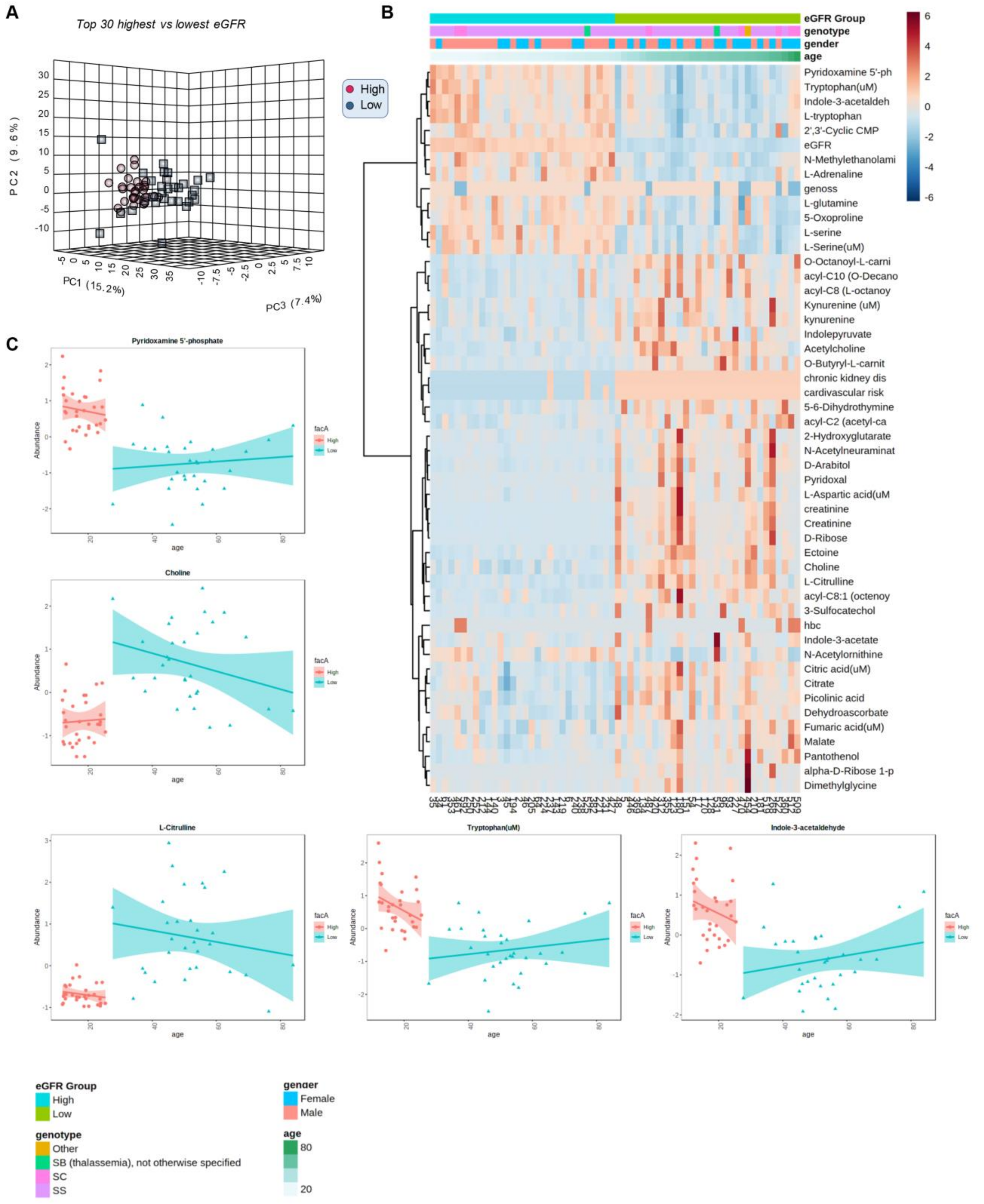

**Supplementary Figure 6** ***Plasma metabolomics of patients enrolled in the WALK-PHASST study with high and low eGFR*** tracks with genotypes (high in SC, low in SS) and shows distinct metabolic phenotypes, as gleaned by principal component analysis (**A**), hierarchical clustering analysis (**B**) and line plots (**C** – x axis tracks with patients' age).

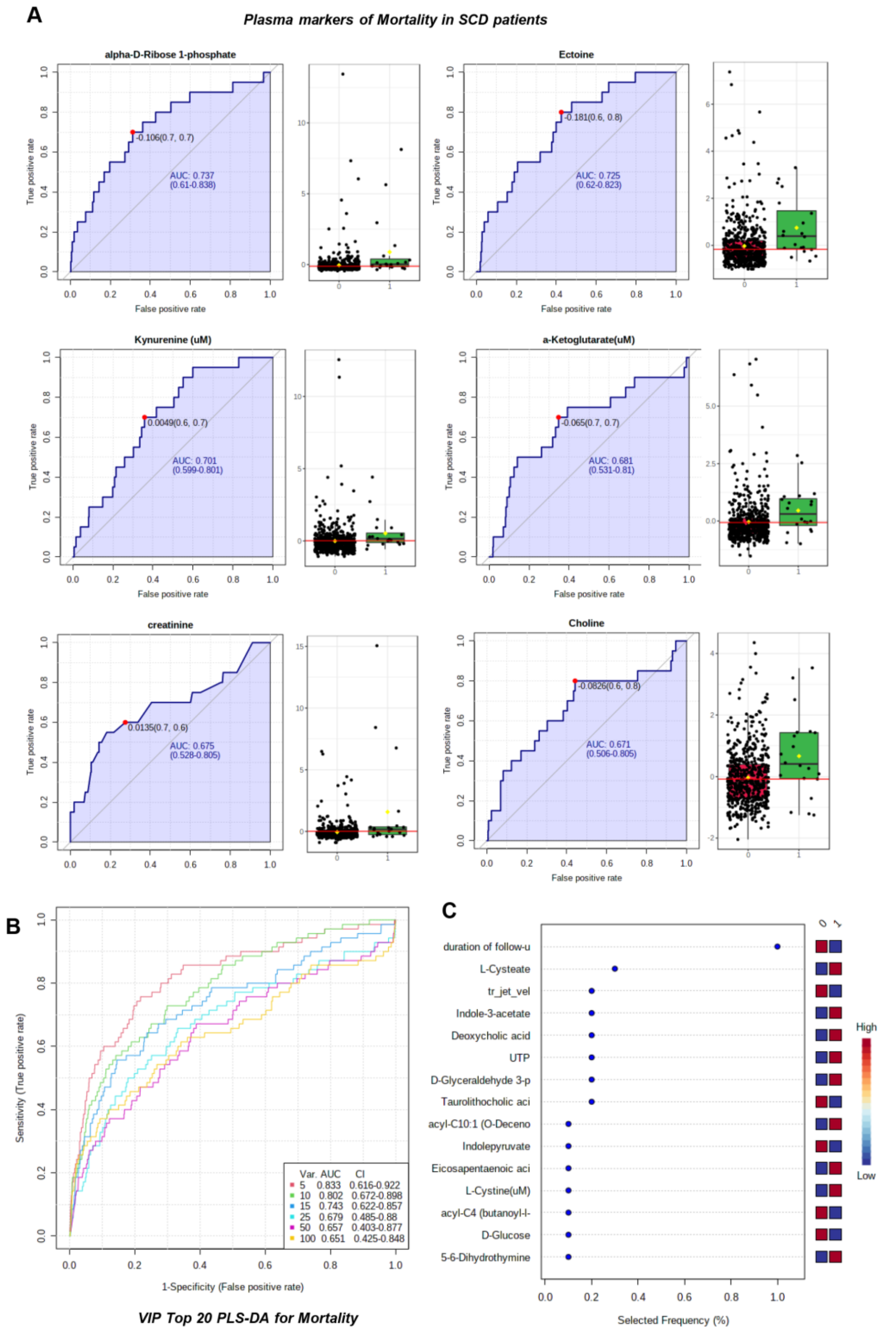

**Supplementary Figure 7** ***Plasma metabolic markers of mortality*** as determined by receiver operating characteristic curves (ROC – **A**), multivariate analyses via random forest (**B**). The top 15 variables from this analysis are reported in **C**.

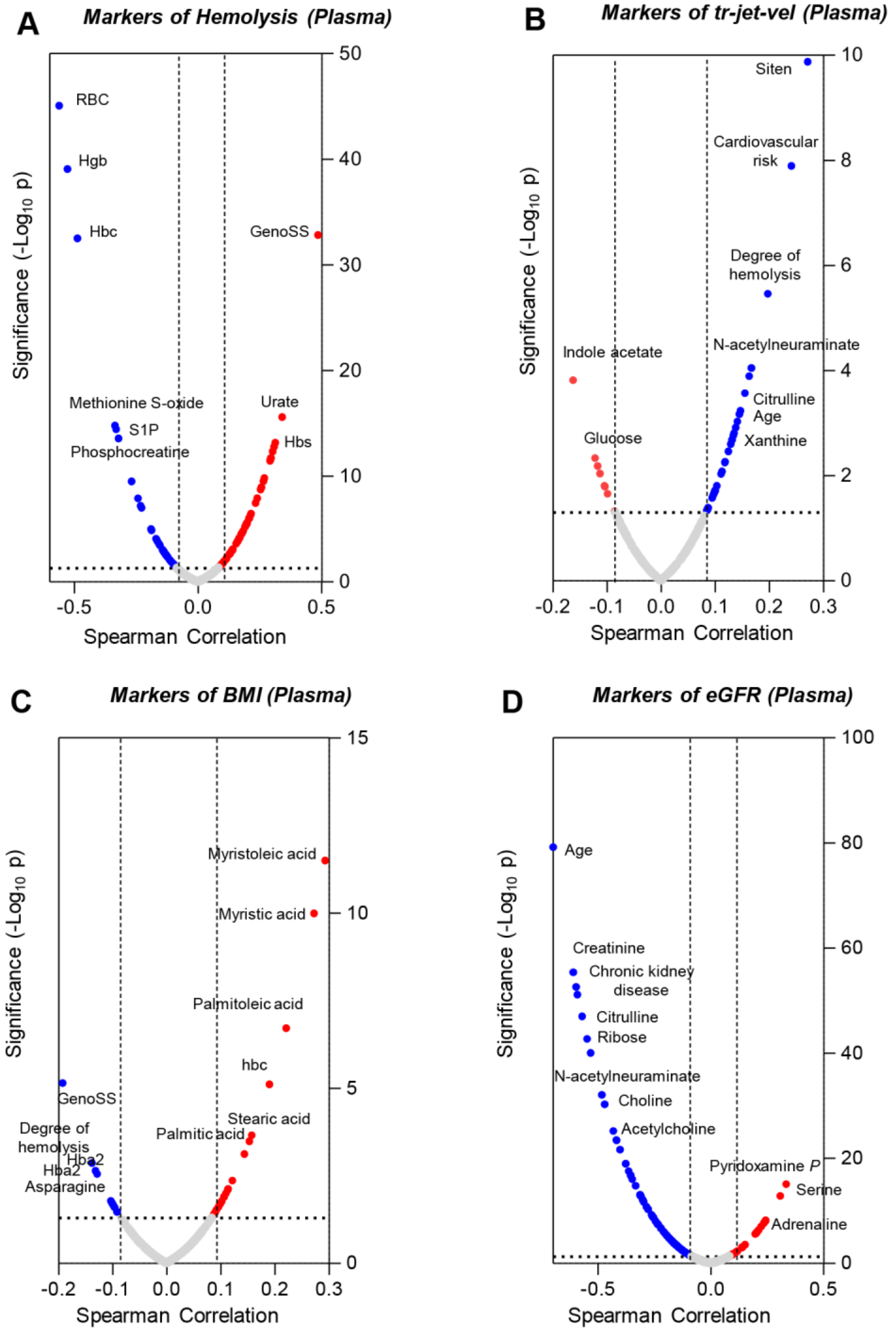

**Supplementary Figure 8** ***Plasma metabolomics correlates to clinical and hematological parameters,*** including the degree of hemolysis (**A**), TRV (**B**), body mass index (BMI – **C**), estimated glomerular filtration rate (eGFR - **D**).

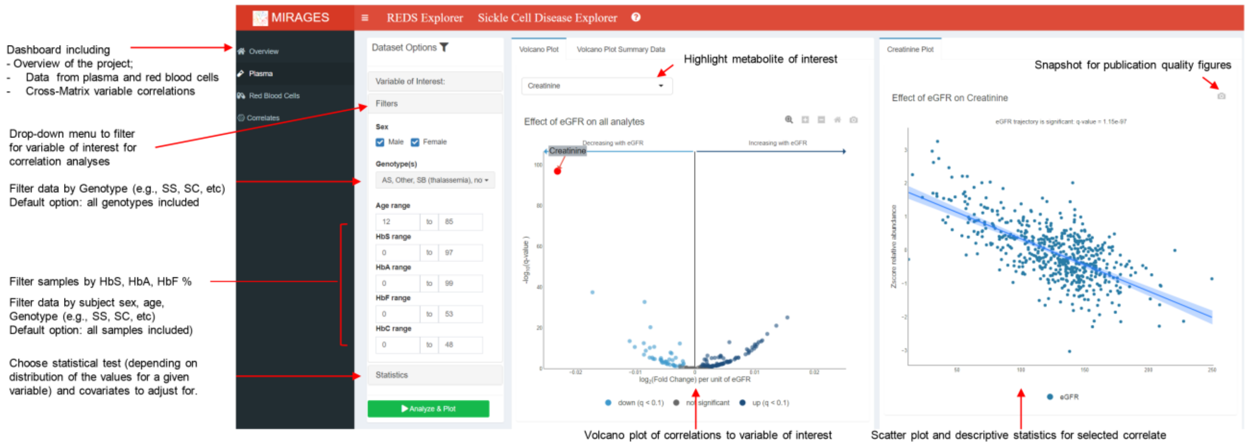

**Supplementary Figure 9** ***Overview of the MIRAGES Sickle Cell Disease portal*** which offers the opportunity to perform streamlined data analysis of plasma, red blood cells or cross-matrix analyses of metabolomics data as a function of relevant clinical or hematological covariates, patients' demographics data. The portal is freely accessible at:
https://mirages.shinyapps.io/SickleCellDisease/

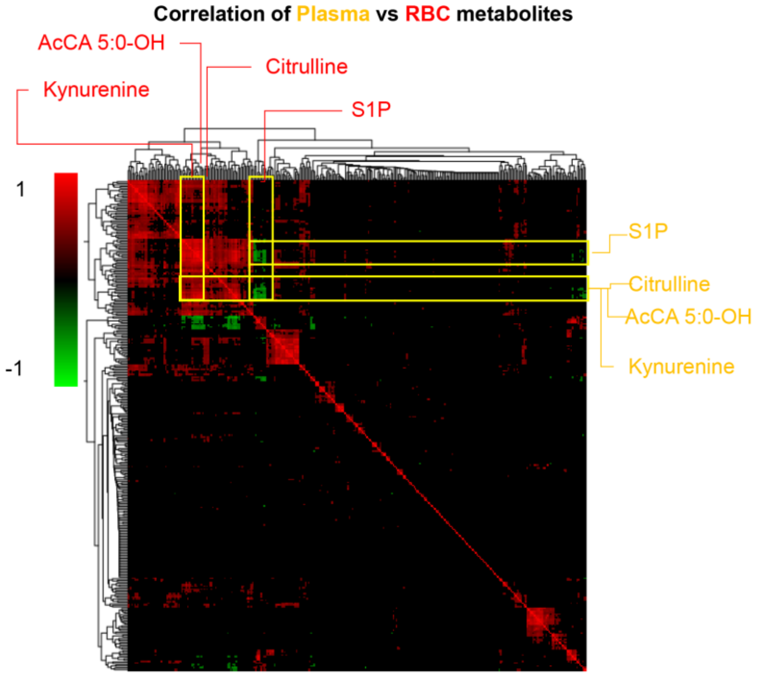

**Supplementary Figure 10** ***Plasma vs RBC metabolites: correlation matrix*** Plasma metabolite levels from the WALK PHASST cohort were matched to RBC metabolomics measurements (*D'Alessandro et al. Am J Hematol 2023*[21]). Results indicated a strong positive cross-matrix correlation for almost all metabolic markers of cardiorenal dysfunction, except for sphingosine 1-phosphate (S1P), whose levels were elevated in SS RBCs and SC plasma, suggesting an increased synthesis or decreased export in the more clinically severe SS genotype

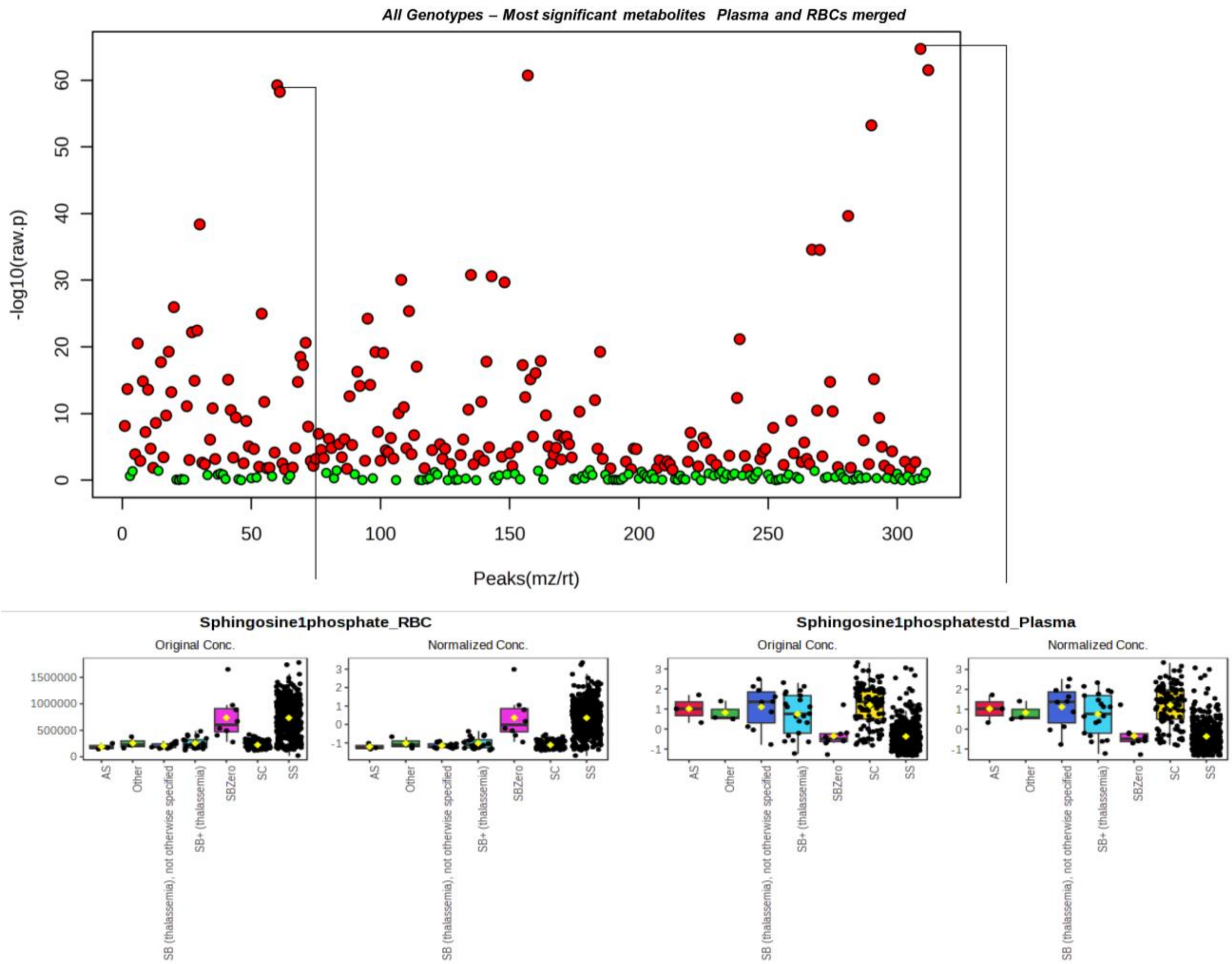

**Supplementary Figure 11** ***ANOVA of merged metabolomics data from plasma and red blood cells from our recent Walk-PHASST study.[11]*** Determination of the most significantly impacted metabolites in plasma (present study) and RBCs[11] from the Walk-PHASST trial are shown in the Manhattan plot at the top of the figure, where the y axis indicates -log10 of p-values by ANOVA based on patients' genotypes and HbS%. Of note, sphingosine 1-phosphate (S1P) in plasma and RBC ranked amongst the top 5 most significantly impacted metabolites of both matrices, though its levels were higher in SS RBCs and lower in SS plasma compared to other genotypes (box plots at the bottom of the figure).
